# A role for the P2Y1 receptor in nonsynaptic cross-depolarization in the rat dorsal root ganglia

**DOI:** 10.1101/702563

**Authors:** Gil B. Carvalho, Yatendra Mulpuri, Antonio Damasio, Igor Spigelman

**Author notes:** Authors for correspondence, Igor Spigelman, PhD, Professor, Division of Oral Biology & Medicine, School of Dentistry, University of California, Los Angeles, 10833 Le Conte Ave., Office: 63-078 CHS, Los Angeles, CA 90095-1668; Antonio Damasio, MD, PhD, Professor, Brain and Creativity Institute, University of Southern California, 3620A McClintock Avenue, Los Angeles, CA 90089-2921, (213) 821 – 2377.

## Abstract

Non-synaptic transmission is pervasive throughout the nervous system. It appears especially prevalent in peripheral ganglia, where non-synaptic interactions between neighboring cell bodies have been described in both physiological and pathological conditions, a phenomenon referred to as cross-depolarization (CD) and thought to play a role in sensory processing and chronic pain. CD has been proposed to be mediated by a chemical agent, but its identity has remained elusive. Here, we report that in the rat dorsal root ganglion (DRG), the P2Y1 purinergic receptor (P2RY1) plays an important role in regulating CD. The effect of P2RY1 is cell-type specific: pharmacological blockade of P2RY1 inhibited CD in A-type neurons while enhancing it in unmyelinated C-type cells. In the nodose ganglion of the vagus, CD requires extracellular calcium in a large percentage of cells. In contrast, we show that in the DRG extracellular calcium appears to play no major role, pointing to a mechanistic difference between the two peripheral ganglia. Furthermore, we show that DRG glial cells also play a cell-type specific role in CD regulation. Fluorocitrate-induced glial inactivation had no effect on A-cells but enhanced CD in C-cells. These findings shed light on the mechanism of CD in the DRG and pave the way for further analysis of non-synaptic neuronal communication in sensory ganglia.

**Highlights:**

- The purinergic receptor P2RY1 plays a regulatory role in non-synaptic cross-depolarization (CD) in the mammalian DRG
- The effect of P2RY1 is cell type-specific: it enhances CD in myelinated A-type neurons, but inhibits it in unmyelinated C-cells
- CD in the DRG does not require extracellular calcium. This is in contrast with the nodose ganglion, where extracellular calcium plays an important role in nonsynaptic interactions
- CD is also modulated by DRG glial cells. Glia selectively inhibit CD in C-type neurons

## Introduction

Although research on neuroscience predominantly focuses on the synaptic modality of information transfer, non-synaptic transmission also represents an important and extremely prevalent phenomenon in the nervous system of mammals, including humans—for example, many of the major neurotransmitters can act non-synaptically (Herkenham 1987, Descarries, Gisiger et al. 1997, Descarries and Mechawar 2000, Vizi, Fekete et al. 2010, Ren, Qin et al. 2011). As opposed to synapses, where diffusion of the neurotransmitter is generally confined to a narrow intercellular gap, in nonsynaptic communication the signaling agent diffuses, relatively unrestrained, in the extracellular space, potentially reaching distant targets (Agnati, Zoli et al. 1995, Carvalho and Damasio 2019). Non-synaptic communication appears particularly prominent in peripheral sensory ganglia (Amir and Devor 1996, Oh and Weinreich 2002, Kim, Anderson et al. 2016, Du, Hao et al. 2017). For example, in dorsal root ganglia (DRG), where synapses are exceedingly rare (Lieberman 1976), most neurons can be depolarized following repetitive activation of neighboring cells (Amir and Devor 1996). This phenomenon, known as cross-depolarization (CD), is predominantly excitatory (Amir and Devor 1996) and can be exacerbated upon injury, but is present in healthy neurons in physiological conditions (Devor and Wall 1990, Amir and Devor 1996). CD has also been described in the nodose ganglion of the vagus (Oh and Weinreich 2002) and may therefore represent a general physiological property of intraganglionic neuronal communication. Despite its prevalence and probable physiological relevance, the exact function of the intraganglionic non-synaptic crosstalk remains mysterious and its mechanisms remain elusive. They may involve the release of diffusible neurochemical substances (Amir and Devor 1996), but this scenario remains unconfirmed and the identity of the molecule(s) involved has not been elucidated.

Our results implicate the P2Y1 purinergic receptor (P2RY1) in non-synaptic CD of DRG neurons. Strikingly, our findings indicate that P2RY1 operates with cell-type specificity. P2RY1 blockade inhibits CD in myelinated A-type neurons, but enhances it in unmyelinated C-cells. We also report that CD mechanisms may not be conserved across peripheral ganglia. Eliminating extracellular calcium had no significant effect on CD magnitude in the DRG, whereas it is essential in a large number of nodose ganglion neurons (Oh and Weinreich 2002). Finally, we report a role for DRG glia in CD. Glial function blockade selectively enhanced CD in C-type neurons. Our results constitute the first description of a neurochemical mechanism for CD in sensory ganglia and suggest possible physiological roles of this widespread phenomenon.

## Materials and Methods

### Tissue preparation

Male Sprague-Dawley rats weighing 300-350 g were anesthetized with isoflurane and oxygen (5:2 for induction, 3:1 for maintenance). Tissues were exposed in layers and L5 DRG were dissected bilaterally together with their respective spinal nerves and dorsal roots and cleaned of surrounding connective tissue. The preparations were allowed to recover for ~2 hrs at room temperature in Ca^2+^-free artificial cerebrospinal fluid (ACSF, 124 mM NaCl, 3.5 mM KCl, 1.25 mM NaH_2_PO_4_, 26 mM NaHCO_3_, 10 mM glucose, 2 mM MgCl_2_) and then transferred to a recording chamber where complete ACSF (containing 2 mM Ca^2+^) was continuously perfused and bath temperature slowly raised to the experimental setting (34.5 °C). In our experience this procedure maximized the viability of the neurons, in particular C-type cells. Solutions were pre-saturated with carbogen (95% O_2_, 5% CO_2_) to ensure adequate oxygenation and a pH of 7.4.

### Electrophysiology

A glass suction electrode was used to stimulate the spinal nerve and another to record compound action potentials (CAPs) from the DR (Fig. 1A, B) as described in detail elsewhere (Matsuka and Spigelman 2004). This served as an indicator of the integrity of sensory axons in individual preparations. Intracellular recordings were conducted using 1.0 mm OD polished-ends borosilicate glass microelectrodes (Model No G100F-4, Warner Instruments, Hamden, CT) pulled to sharpness with the P-87 Flamming/Brown micropipette puller (Sutter Instrument, Novato, CA). Electrodes were filled with 3 M potassium acetate with tip resistances ranging 30-70 MΩ for A-type and 50-120 MΩ for C-type neurons. Electrodes were placed in holders (G23 Instruments, London, UK) connected to the headstage (HS-2) of an amplifier (Axoclamp-2B, Molecular Devices, San Jose, CA). An A/D converter (Digidata 1440A, Molecular Devices) and a computer equipped with pCLAMP 10 software (Molecular Devices) facilitated acquisition (and analysis) of electrophysiological data. Recordings were obtained with the amplifier in Bridge mode with a balanced electrode resistance. The headstage/electrode assembly was advanced with the aid of a Sonnhof Nanostepper (Scientific Precision Instruments GmbH, Oppenheim, Germany). After a neuron was impaled, the spinal nerve was challenged with a range of stimuli of increasing amplitude (0.1-15 mA) and action potential threshold (rheobase) was determined. Cross-depolarization (CD) was recorded in response to subthreshold stimuli, essentially as previously described (Amir and Devor 1996): a 10 s train of stimuli 85% of rheobase (up to a maximum of 10 mA amplitude), 0.2 ms duration (0.5 ms for C-cells) and 100 Hz frequency was used, except where otherwise specified.

**Fig. 1.**
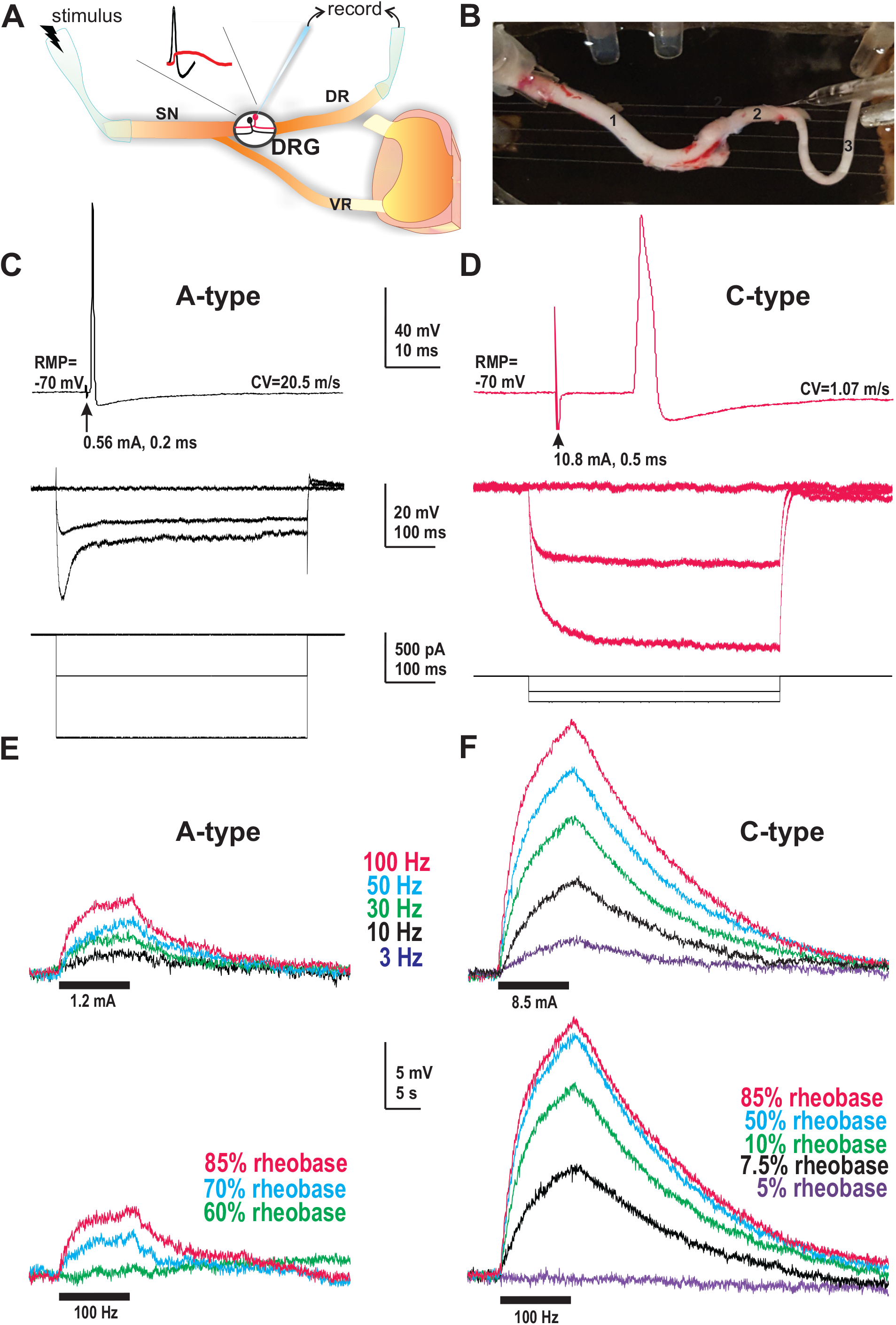
Cross-depolarization (CD) in the rat dorsal root ganglion (DRG). **A**, Schematic of L5 DRG with attached spinal nerve (SN) and dorsal root (DR). Stimulating electrode (left) delivers a stimulus to SN, while intracellular recording electrode (middle) records activity of a single neuronal soma in the DRG. Stimulated sensory fiber (black) experiences an action potential (black trace). Neighboring soma (red) undergoes subthreshold depolarization (red trace) despite the absence of synaptic connections with the stimulated cell. VR, ventral root. **B**, Photograph of the actual experimental setup depicted in (A). VR has been resected. 1, spinal nerve; 2, DRG; 3, DR. **C-F**, Typical electrical properties of neurons analyzed. Left column (**C, E)**, myelinated A-cells. Right column (**D, F**), unmyelinated C-type neurons. **C-D**, Top: Action potential. CV = conduction velocity. RMP = resting membrane potential. Average RMP=-73.3 ± 1.8 mV for A cells (N=22), - 73.9 ± 4.5 mV for C cells (N=10). Arrow indicates stimulus artifact. Bottom: Voltage responses (above) to injection of square hyperpolarizing current pulses (below). R_in_=23.6 MΩ (A, left); 84.9 MΩ (C, right). Average for A cells=19 ± 4 MΩ (N=12). Average for C cells=61.4 ± 13.8 MΩ (N=9). **E-F**, Frequency- and stimulus-dependence of CD voltage responses to SN trains of stimuli. Upper panels illustrate frequency-dependence (black bars, 10 s, 85% of rheobase). Lower panels illustrate stimulus intensity-dependence (black bars, 10 s, 100 Hz). Traces in each panel were obtained sequentially in the same neuron.

A-type and C-type cells were classified based on their electrophysiological properties (Amir and Devor 1996): conduction velocity (CV), the shape of the intracellularly recorded spike (namely the presence (C-type) or absence (A-type) of an inflection on the falling phase) and responses to intracellular hyperpolarizing current injection. CV was calculated by dividing propagation distance (the length of the spinal nerve) by spike latency (time between stimulus and action potential onset) following single suprathreshold stimuli to the spinal nerve. Neurons were discarded if they could not be classified based on the above criteria or if their resting membrane potential (RMP) was less negative than −40 mV (Amir and Devor 1996).

### Determining input resistance

After balancing the amplifier bridge, square wave hyperpolarizing current pulses (1Hz, usually 1–3 nA, 250–500 ms) were delivered intracellularly. Input resistance (R_in_) was calculated from the voltage response using Ohm’s law. To measure input resistance (R_in_) changes during CD, direct intracellular hyperpolarizing current was applied to hyperpolarize the cell to its baseline membrane potential while continuing to deliver the square wave hyperpolarizing current pulses, thus allowing for direct comparison.

### Drug treatment

CD amplitude was determined before and after perfusion of the preparation with pharmacological agents dissolved in ACSF at previously demonstrated effective concentrations. Pharmacological agents and concentrations used: 6-Cyano-7-nitroquinoxaline-2,3-dione [CNQX, 10 μM (Spigelman, Li et al. 2002), Alomone Labs, Jerusalem, Israel], (2R)-2-amino-5-phosphonopentanoic acid [D-AP5, 40 μM (Spigelman, Li et al. 2002), Alomone Labs], MRS2179 tetrasodium salt [100 μM (Agresti, Meomartini et al. 2005); Tocris, Bristol, UK], MRS 2500 tetraammonium salt [1 μM (Grasa, Gil et al. 2009), Tocris] and DL-fluorocitric acid barium salt [100 μM (Chen, Li et al. 2015), Sigma-Aldrich, St. Louis, MO]. The effect of extracellular calcium was tested by perfusing the preparation with calcium-free ACSF balanced with elevated MgCl_2_ (4mM) and containing 100 μM EGTA.

### Data Analysis

Except where otherwise specified, values are shown as average ± SEM. Neurotransmitter receptor antagonist data were analyzed using paired sample t-test (for data normally distributed). Data failing a normality test were analyzed using Wilcoxon signed-rank test. C-cell rundown was corrected for by calculating average rundown experienced by vehicle-treated cells over time (Fig. 2B) and adding that value to the CD magnitude of the ‘post treatment’ group in pharmacological challenges. The effect of fluorocitrate on CD was analyzed using Student’s t-test. Statistical analysis was performed using the SigmaStat 3.5 package (Systat Software, San Jose, CA). P values were considered significant below 0.05.

**Fig. 2.**
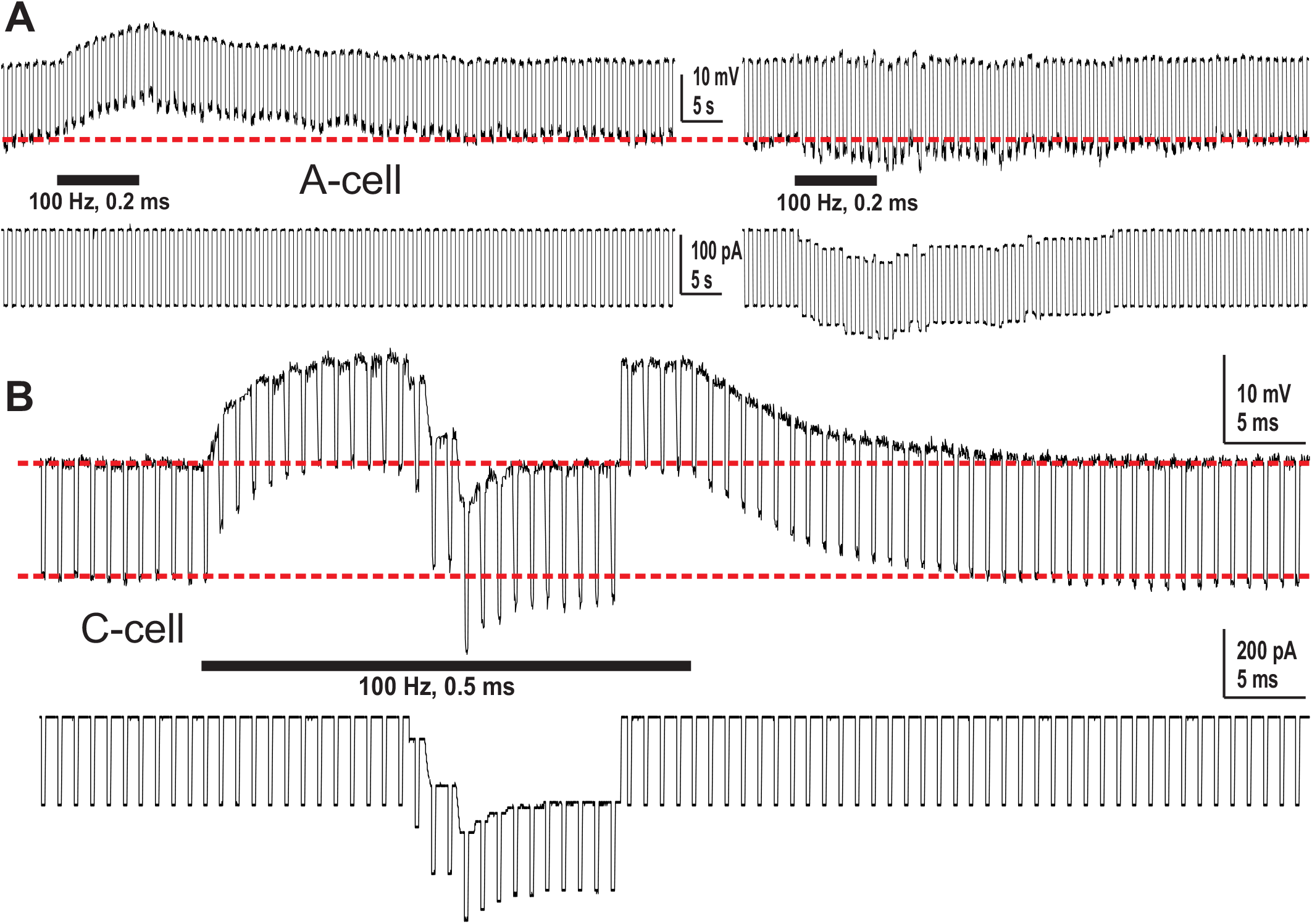
Monitoring input resistance changes during CD. Square wave hyperpolarizing current pulses (1 Hz, usually 1–3 nA, 250-500 ms) were delivered intracellularly. **A**, A-type cell. Left trace shows a CD response to a stimulus train (black bar). Right trace is from the same cell where CD was balanced out using hyperpolarizing direct current, revealing increased voltage responses to the square wave hyperpolarizing current pulses during CD. **B**, C-type cell. Neuron was challenged with a stimulus train (black bar) and CD balanced out by hyperpolarizing direct current. Increased voltage responses to the hyperpolarizing current pulses are seen compared to baseline, indicative of increased input resistance.

## Results

In the DRG, synapses are virtually nonexistent and non-synaptic interactions can be observed in a large number of neurons (Amir and Devor 1996). To study this phenomenon, L5 DRG attached to the spinal nerve (SN) and dorsal root (DR) were excised and kept in near physiological conditions (Fig. 1A-B), allowing the SN to be challenged with electrical stimuli while intracellular recordings are obtained from individual neuronal cell bodies in the ganglion (Fig. 1C-F). SN were stimulated as previously described (Amir and Devor 1996): stimuli were set at 85% of intensity required to elicit an action potential (rheobase), up to a maximum of 10 mA, ensuring stimuli were sub-threshold for the particular neuron being studied.

Consistent with previous reports, cells typically responded to subthreshold repetitive axonal stimuli with a gradual depolarization, sometimes reaching a plateau, with a gradual repolarization following stimulus cessation (average CD magnitude: A cells=1.9 ± 0.6mV, N=24; C cells=4.2 ± 1.2mV, N=13) (Fig. 1E-F). CD response was both stimulus frequency- and intensity-dependent (Fig. 1E-F) and was associated with an increase in input resistance (average increase during CD in A cells=24.8 ± 8.6%, p=0.009, N=8; C cells=13 ± 5%, p=0.034, N=12, Fig. 2), as previously reported (Amir and Devor 1996).

We next set out to elucidate the mechanism of CD in the DRG, which has been hypothesized to involve a neurochemical agent (Amir and Devor 1996). In order to identify neurotransmitter(s) involved in CD regulation, we conducted a pharmacological screen. DRG were challenged with commercially available neurotransmitter receptor antagonists and CD recorded before and after drug treatment. In control conditions (perfusion with ACSF, in the absence of pharmacologic agents), magnitude of CD in A-type cells did not change significantly after 8-10 min, the time window used for drug perfusion (p=0.9, N=12, Fig. 3A). In contrast, unmyelinated C-cells exhibited a significant decline in CD amplitude over time (rundown observed in 44% of cells, average decline=17%, p=0.015, N=9, Fig. 3B). C-type neurons were generally more challenging to record from, and less stable, than their A-type counterparts, which may explain this occurrence. Therefore, subsequent pharmacology data obtained from C-cells were normalized to factor out this treatment-independent rundown (see Methods section).

**Fig. 3.**
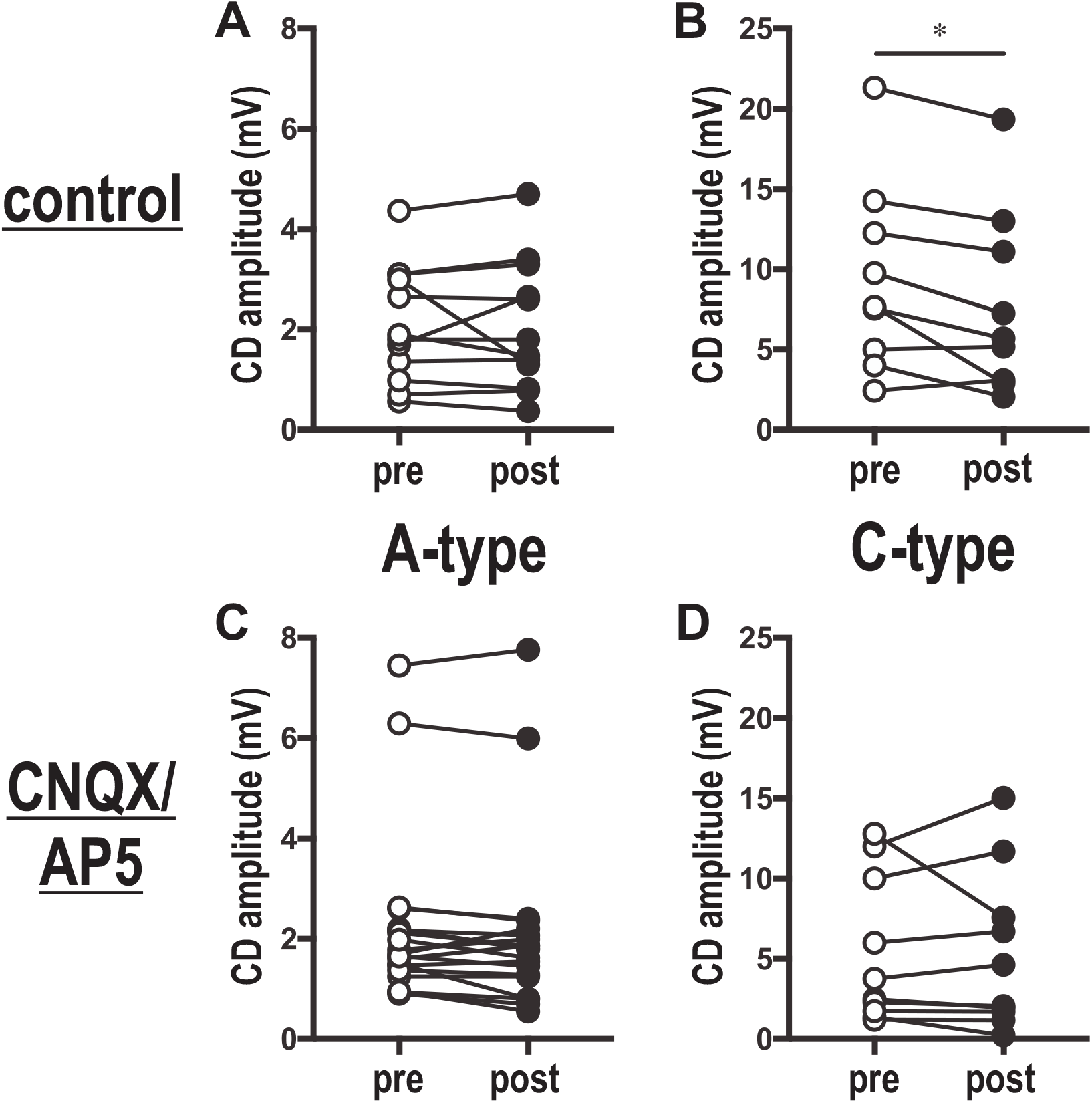
Effect of ionotropic glutamate receptor blockade on CD in the DRG. **A-B**, ACSF perfusion (control). CD recorded before and after 8-10 min perfusion. **A**, Magnitude of CD does not change significantly with time in A-type cells (p=0.9, N=12). **B**, Magnitude of CD decreases over time in unmyelinated C-cells (average rundown=17%, p = 0.015, N=9). **C**, Simultaneous treatment with AMPAR/KAR/NMDAR antagonists (10 μM CNQX + 40 μM AP5) does not significantly impact CD magnitude in A-type neurons. Amplitude of CD response plotted for each individual neuron in the presence (“post”) and absence (“pre”) of drug treatment. p = 0.056, N = 20. **D**, AMPAR/KAR/NMDAR blockade does not significantly impact CD magnitude in C-type neurons. Values were normalized to account for the rundown observed in control conditions (see text). p = 0.9, N = 10. **C-D,** “pre” and “post” drug measurements were 8-10 mins apart.

Given the fact that virtually all DRG neurons are glutamatergic (Huettner, Kerchner et al. 2002), we first tested selective blockers of α-amino-3-hydroxy-5-methyl-4-isoxazolepropionic acid (AMPA), kainate and N-methyl-D-aspartate (NMDA) glutamate receptors. Combined application of CNQX (10 μM) and AP5 (40 μM) had no statistically significant effect on CD in A-type (p = 0.056, N=20, Fig. 3C) or C-type (p=0.9, N=10, Fig. 3D) neurons. These results do not completely exclude an effect of glutamate in a small subset of cells—in both cell types there was a slight nonsignificant decrease in average CD magnitude with CNQX/AP5 application (A-type: average CD went from 2.24mV to 2.1mV; C-type: from 5.365mV to 5.276mV)—but they strongly indicate that ionotropic glutamate receptors do not play a major role in CD in the majority of DRG neurons.

We next considered the involvement of purinergic receptors. Adenosine triphosphate (ATP) is released from both peripheral and central sensory neuron terminals (Bodin and Burnstock 2001, Sawynok and Liu 2003), as well as within sensory ganglia (Matsuka, Neubert et al. 2001, Matsuka, Ono et al. 2008), and is subsequently hydrolyzed to ADP and further to AMP and adenosine by ectonucleotidases in the DRG (Vongtau, Lavoie et al. 2011). We did not focus our analysis on the ionotropic P2X2 and P2X3 receptors because P2X receptors have been shown to mediate membrane potential depolarization associated with increases in membrane conductance (Hiruma and Bourque 1995, Burgard, Niforatos et al. 1999, Grubb and Evans 1999, Xu and Huang 2002), in contrast to the decrease associated with CD (Amir and Devor 1996). Moreover, P2X2 and P2X3 receptors are generally subject to relatively rapid desensitization (Brändle, Spielmanns et al. 1997, Sokolova, Skorinkin et al. 2006), whereas CD is a non-desensitizing phenomenon up to at least 10 seconds of duration (Amir and Devor 1996) (Fig. 1). Thus, we next tested the involvement of P2RY1, a purinergic G-protein coupled receptor activated predominantly by adenosine diphosphate (ADP) and which has been associated with increased input resistance and blocking of ion channels (Filippov, Brown et al. 2000, Filippov, Choi et al. 2006). P2RY1 is expressed in sensory neurons and has been implicated in thermosensation and nociception (Malin and Molliver 2010, Molliver, Rau et al. 2011, Jankowski, Rau et al. 2012). Treatment with the P2RY1 antagonist, MRS 2179, significantly reduced CD magnitude in A-type cells (p=0.007, N=16, Fig. 4A, E). A dampened CD response was observed in ~44% of A-type neurons (average reduction=46.5 ± 9%; 50% of cells tested showed no response and 6% showed increased response).

**Fig. 4.**
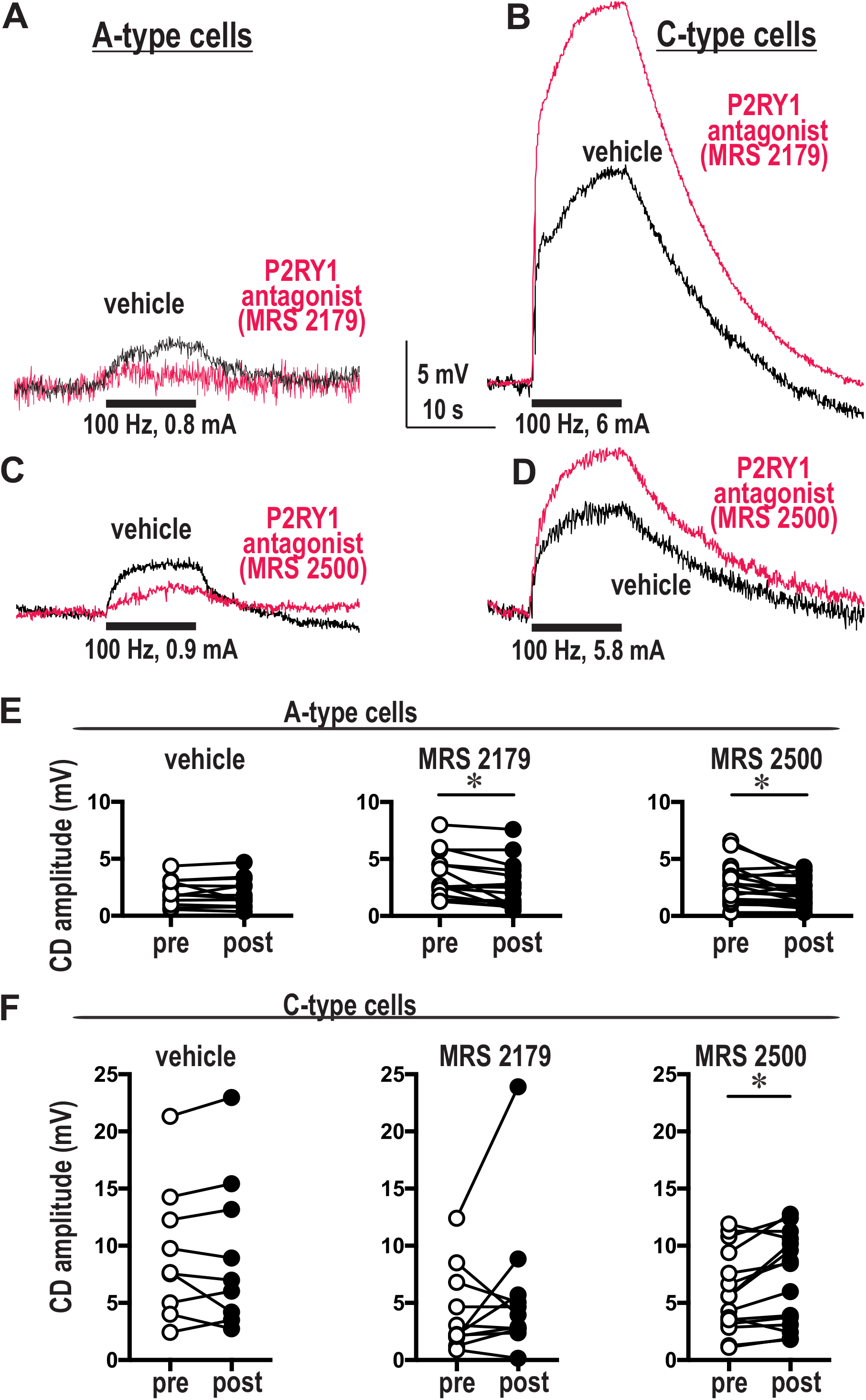
The purinergic receptor P2RY1 regulates CD in the DRG. Treatment with either of two different P2RY1 antagonists (100 μM MRS 2179 or 1 μM MRS 2500) hinders CD in A-type neurons (**A, C, E**) while enhancing it in C-type cells (**B, D, F**). **A-D**, Voltage recordings upon application of stimulus trains (represented by black bars) in the presence and absence of P2RY1 antagonists. Example traces from responder neurons show CD in control conditions (artificial cerebrospinal fluid alone, black traces) and altered response in the presence of antagonist drug (red traces). Left panels (**A**, **C**), A-type cells; Right panels (**B**, **D**), C-type cells; Top panels (**A**, **B**), MRS 2179; Bottom panels (**C**, **D**), MRS 2500. **E-F**, Amplitude of CD response plotted for each individual neuron in the presence and absence of drug treatment. **E**, A-type cells. Vehicle, N=12. MRS 2179, p=0.007, N=16. MRS 2500, p=0.012, N=19. **F**, C-type cells. Vehicle, N=9. MRS 2179, p=0.27, N=11. MRS 2500, p=0.018, N=15. C-type neuron values (**F**) were normalized to account for the rundown observed in control conditions (see text). *, p<0.05, Wilcoxon signed-rank test (**E**) or paired t test (**F**).

To confirm our observations, we used an additional P2RY1 blocker. MRS 2500 tetraammonium salt is a highly potent, selective and stable antagonist of P2RY1 (Hechler, Nonne et al. 2006)—it has 100-fold higher binding affinity than MRS 2179 (Hechler, Nonne et al. 2006) and superior potency (Jiménez, Clavé et al. 2014) [K_i_=0.78nM; IC_50_=8.4nM (Kim, Ohno et al. 2003), whereas the reported IC_50_ for MRS 2179 is one order of magnitude higher, in the 100-300nM range (Kaiser and Buxton 2002)]. The effect of MRS 2500 treatment was similar to MRS 2179. A significant reduction of CD magnitude was seen in A-type cells (p = 0.012, N=19, Fig. 4C, E), with ~47% of cells showing a dampened CD response (average reduction=41 ± 3%; 42% of cells tested showed no change and 10.5% showed increased response). These data suggest that P2RY1 blockade significantly inhibits CD in a substantial subset of A-type neurons.

We next asked if P2RY1 plays a similar role in unmyelinated, C-type cells. Both P2RY1 pharmacological inhibitors showed a trend toward enhancing CD in a subset of C neurons, with ~46% of cells showing an increased response with either drug (MRS 2179: average increase=150 ± 50%, 27% of cells showed no change and 27% showed decreased CD; MRS 2500: average increase=53 ± 8%, ~46% of cells showed no change and 7% showed decreased CD, Fig. 4B, D, F). This change reached statistical significance for the potent, selective antagonist MRS 2500 (p = 0.018, N=15; MRS 2179: p=0.27, N=11, Fig. 4F).

Taken collectively, our observations suggest that P2RY1 plays a regulatory role in the DRG, with a cell type-specific effect—promoting CD in large myelinated A-type neurons while inhibiting it in unmyelinated C-type cells.

We asked if the effect of P2RY1 blockade on CD magnitude can be explained by changes in input resistance. For this test, we focused on the potent, selective MRS 2500. We did not find a significant correlation between input resistance change (ΔR_in_) and CD change (ΔCD) with MRS 2500 application (A cells, p=0.098, N=7; C cells, p=0.642, N=8, Pearson correlation). However, given the modest magnitude of the changes in input resistance taking place during CD (~25% for A cells, ~13% for C cells, Fig. 2), and the fact that drug-induced changes involve a much larger timescale than electrical stimuli (8-10 minutes vs a few seconds), we cannot rule out changes occurring below detection threshold.

We also did not find a consistent association between changes in CD magnitude with P2RY1 blockade (ΔCD) and cellular electrical properties. In A-type neurons, ΔCD was not significantly correlated with baseline CD magnitude (p=0.75 for MRS 2179 / p=0.11 for MRS 2500, Pearson correlation), baseline input resistance (p=0.18 / p=0.166), resting membrane potential (p=0.098 / p=0.88) or rheobase (p=0.15 / p=0.66). In C cells, baseline CD (p=0.6 / p=0.255) and input resistance (p=0.65 / p=0.86) also showed no correlation with ΔCD. Resting membrane potential and rheobase were statistically associated with ΔCD upon MRS 2500 treatment (RMP: *r*=0.7, p=0.00244, N=15; rheobase: r=-0.554, p=0.04, N=14), but these associations were not recapitulated with MRS 2179 (RMP: p=0.93; rheobase: p=0.32). In sum, these results do not clearly establish unique cellular properties of neurons responding to P2RY1 blockade.

CD has also been reported in the nodose ganglion of the vagus, where extracellular calcium plays a central regulatory role—eliminating extracellular calcium abrogates the CD response in 60% of neurons (Oh and Weinreich 2002). We asked if a similar mechanism exists in the DRG. Strikingly, we found no significant decrease of CD magnitude when extracellular calcium was removed, irrespective of cell type (A-type neurons: p=0.744, N=16; C-type neurons: p=0.13, N=10, Fig. 5). This observation suggests that CD may involve different cellular mechanisms in different peripheral ganglia.

**Fig. 5.**
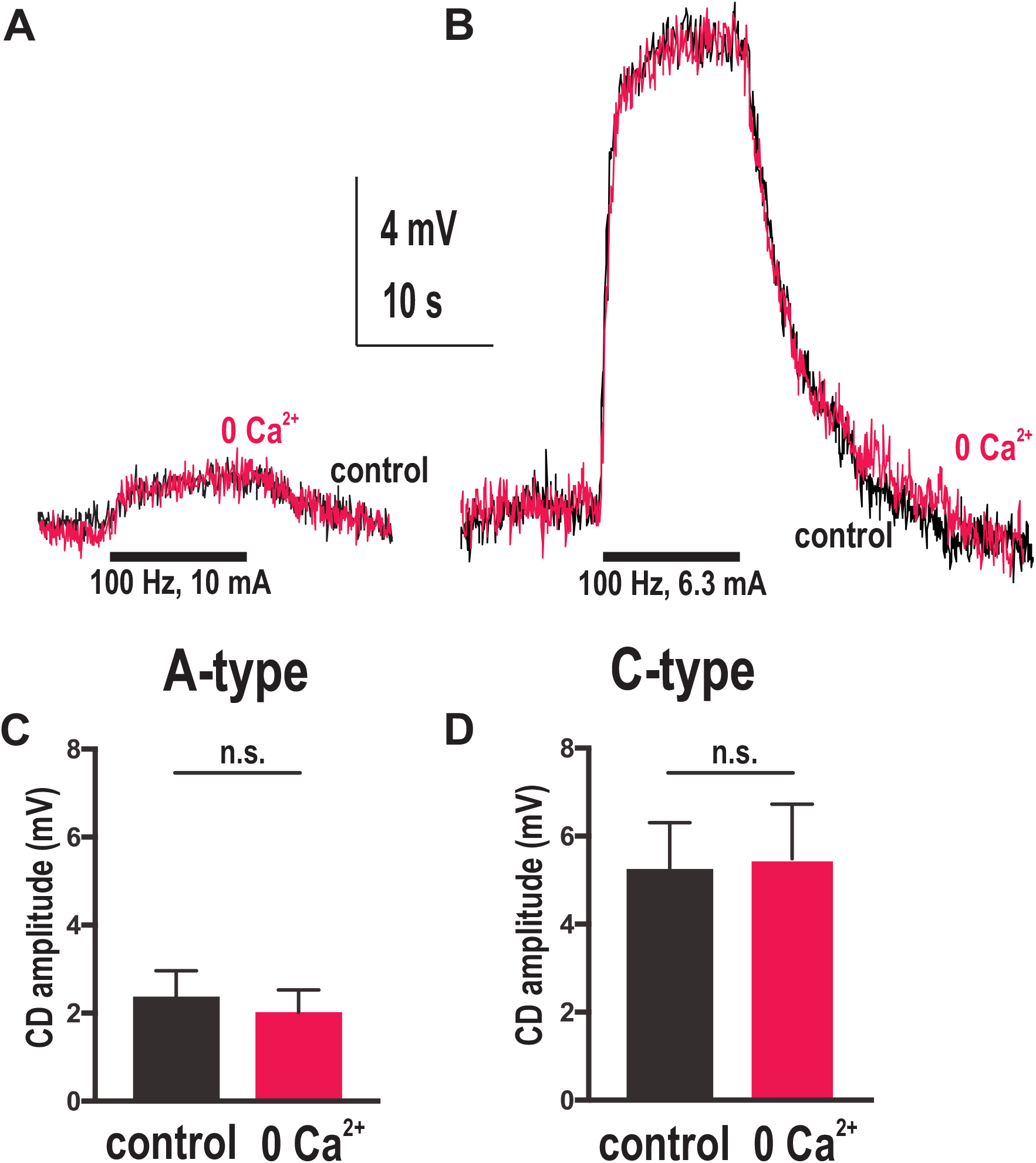
Extracellular calcium does not significantly affect CD magnitude. Left panels (**A**, **C**), A-type cells; Right panels (**B**, **D**), C-type cells. **A-B**, Voltage recordings upon application of stimulus trains (represented by black bars) in the presence and absence of extracellular calcium. Example traces show CD in control conditions (regular artificial cerebrospinal fluid, black traces) and zero (0) extracellular calcium (red traces). **C-D**, Average CD magnitude in the presence and absence of extracellular calcium. **C**, A-type neurons, p=0.744, N=16. **D**, C-type neurons: p=0.13, N=10.

Glia have been implicated in nonsynaptic neuronal communication (Syková and Chvátal 2000). In the DRG, satellite glial cells (SGCs) surround neuronal cell bodies and participate in glia/neuron paracrine interactions (Chen, Zhang et al. 2008, Rozanski, Li et al. 2013). We asked if glia also play a role in CD. SGC metabolism can be selectively blocked without affecting neuronal function using the gliotoxin fluorocitrate (FC) (Chen, Zhang et al. 2008). Since the effect of FC is irreversible and requires a lengthy incubation (90 min), rather than analyzing individual cells before and after treatment we focused on changes in average CD magnitude. Average CD magnitude in A-cells was not significantly affected by glial blockade (mean CD=1.9 ± 0.6mV in control conditions; 1.9 ± 0.59mV in the presence of FC; p=0.93; N=24 for control, 26 for FC, Fig. 6A). However, in C-cells average CD was significantly elevated upon FC-induced glial blockade (mean CD=3.3 ± 1.2mV in control conditions; 10 ± 3mV in the presence of FC; p=0.043; N=10 for control, 8 for FC, Fig. 6A). FC incubation did not significantly affect the A-cell compound action potential (p=0.7, N=6, Fig. 6B), suggesting the increased response seen in C-type neurons is not merely a consequence of a larger population of A-type neurons firing. These observations indicate that SGCs play a cell type-specific role in regulating CD in the DRG, inhibiting nonsynaptic crosstalk specifically in unmyelinated C-type neurons.

**Fig. 6.**
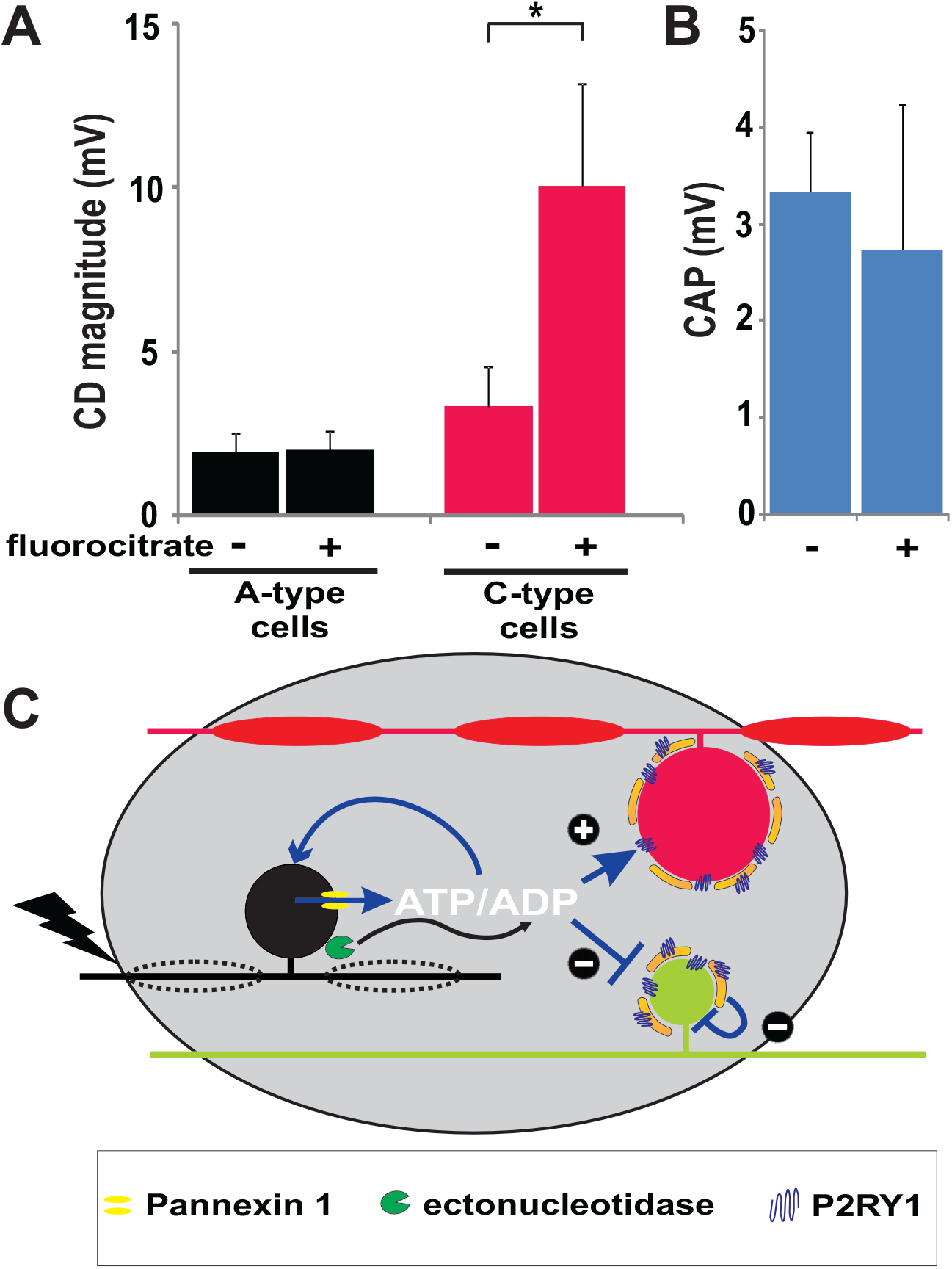
The role of glia in CD in the DRG. **A,** The DRG satellite glial cells (SGCs) mediate CD in C-, but not A-type neurons. The average CD magnitude of neurons in whole-mount ganglia in control conditions was compared to average CD magnitude in the same ganglia after incubation with 100 μM fluorocitrate (FC), a selective inhibitor of glial metabolism. The treatment showed no significant effect in A-cells (p=0.93), but strongly enhanced CD magnitude in C-type neurons (mean CD=3.3 ± 1.2mV in control conditions; 10 ± 3mV in the presence of FC; p=0.043). Bars represent mean ± SEM. *, p<0.05, Student’s t-test. Plus sign (+), glial function intact, ACSF perfusion (control condition); Minus sign (-), glial function impaired, FC perfusion (experimental condition). N=24 (A, control), 26 (A, FC-treated), 10 (C, control), 8 (C, FC-treated). **B**, FC treatment does not significantly impact A-cell compound action potential (CAP) magnitude. p = 0.7, Student’s t test, N=6. **C**, Mechanistic model for CD in the DRG. Stimulated fiber (black) experiences an action potential, triggering ATP release (mediated by channels such as Pannexin 1) into the surrounding extracellular space. Purines (primarily ADP resulting from the extracellular hydrolysis of ATP by ectonucleotidases) bind to P2RY1 on the membrane of neighboring A-cells (red), inducing CD (plus sign). In unmyelinated C-cells (green), on the other hand, the purines/P2RY1 pathway inhibits CD, with SGCs also playing a regulatory role (T-ended arrows and minus signs). Whether purine-induced CD inhibition in C-neurons is mediated by glial and/or neuronal P2RY1 is presently unclear.

## Discussion

Our findings implicate the purinergic P2RY1 receptor in CD in the rat DRG: P2RY1 blockade inhibited CD in myelinated A-type cells, but enhanced it in C-cells. These results constitute the first report of a neurochemical mechanism for CD in sensory ganglia.

CD is almost universal among DRG neurons (Amir and Devor 1996)—a region of the nervous system where synapses are exceedingly rare. It also seems prevalent in other sensory ganglia (Oh and Weinreich 2002), and may thus be a general property of the peripheral nervous system. CD has been shown to be predominantly excitatory, modulating the probability of activation in the affected sensory neurons (Amir and Devor 1996), although cross-inhibition of mechanoreceptive inputs in DRG of cats after peripheral inflammation has also been demonstrated (Xu and Zhao 2003). CD has been proposed to be neurotransmitter-mediated, but the specific molecule(s) at stake have remained mysterious. DRG neurons are primarily glutamatergic (Huettner, Kerchner et al. 2002), and both anatomical and functional studies show widespread ionotropic glutamate receptor expression in the DRG (Shigemoto, Ohishi et al. 1992, Coggeshall and Carlton 1997, Lee, Bardoni et al. 2002, Bardoni, Torsney et al. 2004, Li, McRoberts et al. 2004). Nonetheless, we found that ionotropic glutamate receptors (AMPA, kainate and NMDA) do not appear to play a major role in CD (Fig. 3). Simultaneous pharmacological blockade using CNQX/AP5 did not significantly affect CD magnitude, although a modest trend was observed (Fig. 3). Thus, although glutamate may play a role in CD in a subset of neurons, it does not appear to substantially regulate CD in most of the DRG. In contrast, the purinergic P2RY1 appears to play a significant effect in substantial subsets of both A- and C-type cells, albeit with cell type-specific effects, namely promoting CD in large myelinated (A-type) and inhibiting it in small, unmyelinated (C-type) neurons.

The role of P2RY1 in CD reported here is consistent with previous reports. CD involves decreased membrane conductance (Amir and Devor 1996) and P2RY1 can trigger excitation of DRG neurons via an inhibition of Kv7 potassium channels (Yousuf, Klinger et al. 2011, King and Scherer 2012, Gafar, Rodriguez et al. 2016). This downstream mechanism may at least partly explain the regulatory role of P2RY1 in CD. Our observations are also in keeping with the role of purines in intercellular communication in sensory ganglia (Gerevich, Müller et al. 2005, Magni and Ceruti 2013, Magni, Merli et al. 2015, Rajasekhar, Poole et al. 2015, Warwick and Hanani 2016).

Our findings allow us to formulate a first, tentative model of CD in the DRG (Fig. 6C). Purines released as a result of action potentials (Matsuka, Neubert et al. 2001, Matsuka, Ono et al. 2008) diffuse through the ganglionic stroma and modulate P2RY1 on neighboring cells, with cell-type specific consequences—P2RY1-dependent depolarization of myelinated A-cells and inhibition of CD in C-cells. Our findings also identify an SGC-mediated inhibition of CD in C-type neurons. Whether this accounts for the effect of P2RY1 in C cells or constitutes a separate, additional mechanism is presently unclear. In the DRG, P2RY1 is expressed in both neurons and glia (Chen, Zhang et al. 2008, Huang, Gu et al. 2013, Magni and Ceruti 2013, Rajasekhar, Poole et al. 2015). Our results indicate that, at least for A-type cells, glial P2RY1 does not play a substantial role in CD, since SGC inactivation does not have a significant impact (Fig. 6A). Thus in large, myelinated A-cells, P2RY1-mediated crosstalk likely represents a direct, neuron-to-neuron nonsynaptic communication pathway. In C-cells, on the other hand, SGCs do appear to modulate CD. These observations are consistent with structural data indicating that molecules in the DRG extracellular space can more easily access large cells with myelinated axons, whereas SGCs form a tighter, less permeable barrier around small C-type neurons (Shinder and Devor 1994). Thus, extracellular purines released by activated neurons may diffuse and interact directly with neighboring A-cells (via P2RY1 expressed on their surface), whereas direct access to C-cells may be hindered by the tighter wrapping of SGCs (Shinder and Devor 1994), with diffusing purines interacting instead with glial P2RY1, and the affected glia in turn regulating CD in the associated small C-type neurons. Alternatively, the effects of glia and P2RY1 identified here may represent separate cellular mechanisms. Additional future work will be required to distinguish between these scenarios. It will also be interesting to probe whether the cell type-specific effect of glia—modulating CD in C-, but not A-cells—explains our observation that P2RY1 antagonists have opposite effects in A- and C-type neurons, blocking CD in the former while potentiating it in the latter. Alternatively, intracellular downstream effectors may execute different action programs in A- and C-neurons, leading to particular cellular responses. Also, more than one ligand may be involved, since P2RY1 is known to bind multiple purines (e.g. ATP and ADP) and to possess at least two discrete extracellular binding sites (Zhang, Gao et al. 2015).

Myelinated and unmyelinated neurons typically exhibit differences in nonsynaptic communication, but those discrepancies are generally localized to axonal projections— such as ephaptic conduction, which tends to occur between unmyelinated, tightly packed fibers (Bokil, Laaris et al. 2001)—presumably a product of myelin acting as a physical barrier to the release and access of diffusing ligands, such as neurotransmitters or other signaling molecules (Syková and Chvátal 2000, Sykova 2004, Sykova 2005, Damasio and Carvalho 2013). To our knowledge, this is the first evidence that myelinated and unmyelinated neurons also differ in the nonsynaptic exchanges taking place intraganglionically, away from nerves proper. Exploring further the relationship between glia and nonsynaptic interactions may shed further light on the neurophysiology of sensory signal processing in the peripheral nervous system.

Our findings indicate that CD regulation is multifactorial. Both A- and C-type neurons were found where P2RY1 pharmacologic modulation showed no clear effect. Moreover, the inhibitory effect of CD by P2RY1 antagonists averaged 41-44% (depending on the compound used). We employed high concentrations of P2RY1 antagonists. For example, for MRS 2179, the experimental concentration used (100 μM) is 30-100x the reported IC50 (Anita Jagroop, Burnstock et al. 2003, Jiménez, Clavé et al. 2014), and MRS 2179 has been shown to be effective even at 10 μM (Baurand, Raboisson et al. 2001). Similarly, for MRS 2500, the experimental concentration (1μM) is ~20x the IC50 (Jiménez, Clavé et al. 2014). Yet neither compound induced a 100% abrogation of CD. These findings indicate that additional regulatory mechanisms for CD likely exist. This scenario is consistent with previous reports suggesting a role for extracellular K^+^, possibly via Nernstian depolarization (Amir and Devor 1996).

We also found that eliminating extracellular calcium did not significantly affect CD magnitude (Fig. 5). CD in the DRG may rely on intracellular calcium stores or be entirely calcium-independent. The large-pore membrane channel Pannexin-1 (Panx1) is involved in ATP release and can be opened by cellular depolarization (Locovei, Wang et al. 2006) yet operates independently from extracellular calcium (Bruzzone, Barbe et al. 2005). Panx1 is expressed in DRG neurons, where it plays a role in neuropathic pain (Zhang, Laumet et al. 2015). CD may involve Panx1-mediated ATP release, which would explain the observed lack of requirement of extracellular calcium. Since extracellular calcium plays a central role in CD in the nodose ganglion of the vagus (Oh and Weinreich 2002), our findings suggest a fundamental mechanistic difference in CD regulation between these two peripheral sensory ganglia.

The physiological role of CD in sensory ganglia is presently unclear. It has been proposed to represent a feedback mechanism, perhaps modulatory of both neuronal metabolism and excitability (Devor 1999). The DRG functions as a gateway for action potential propagation (Krames 2014). Specifically, the somatic/perisomatic compartment—the part of the DRG neuron comprising the cell body, T-junction and the initial axon segment connecting them—is able to modulate the conduction of action potentials from the periphery to the spinal cord, effectively regulating the transmission of sensory information to the CNS (Du, Hao et al. 2014). As the T-junction is the point of lowest safety for action potential propagation in the peripheral sensory pathway, small changes in membrane voltage can determine the success or failure of impulse conduction, particularly in C-cells. Thus, even modest changes in membrane potential at the somatic/perisomatic compartment—as small as 2-4 mV, i.e. well within the range observed in our experiments—have been shown to interfere with fiber excitability, block action potential propagation and significantly affect pain behaviors (Du, Hao et al. 2014). Thus, CD has the potential to modulate sensory information transmission, nociceptive and otherwise. The remarkable cell-specificity described here—P2RY1 shows contrary effects on A- and C-type neurons—suggests a novel process of neural coding between cognate fibers with a putative role in coordinating sensory information transmission and processing. As such, the potential of CD as a productive target for novel pharmacological approaches to pain control is considerable.

CD can be observed in most neurons in the DRG in normal conditions (Amir and Devor 1996), and is exacerbated upon injury (Devor and Wall 1990). CD has also been described in the nodose ganglion of the vagus nerve (Oh and Weinreich 2002), and may thus represent a general feature of peripheral sensory ganglia. Elucidating the physiological role(s) of CD will likely contribute to our understanding of sensory perception and nociception and may also shed light on the mechanisms of interoceptive processing and affect. The first description of a neurotransmitter mechanism for CD in the DRG constitutes a fundamental first step in that direction.

## Acknowledgements

This work was supported by a grant to A.D. from The Mathers Foundation. We thank our colleagues H. Damasio, J. Kaplan, K. Man and M. Henning for insightful discussions and comments on the manuscript.

## References

Agnati, L., M. Zoli, I. Strömberg and K. Fuxe (1995). “Intercellular communication in the brain: wiring versus volume transmission.” Neuroscience 69(3): 711–726.

Agresti, C., M. Meomartini, S. Amadio, E. Ambrosini, B. Serafini, L. Franchini, C. Volonte, F. Aloisi and S. Visentin (2005). “Metabotropic P2 receptor activation regulates oligodendrocyte progenitor migration and development.” Glia 50(2): 132–144.

Amir, R. and M. Devor (1996). “Chemically mediated cross-excitation in rat dorsal root ganglia.” Journal of Neuroscience 16(15): 4733–4741.

Anita Jagroop, I., G. Burnstock and D. P. Mikhailidis (2003). “Both the ADP receptors P2Y 1 and P2Y 12, play a role in controlling shape change in human platelets.” Platelets 14(1): 15–20.

Bardoni, R., C. Torsney, C.-K. Tong, M. Prandini and A. B. MacDermott (2004). “Presynaptic NMDA receptors modulate glutamate release from primary sensory neurons in rat spinal cord dorsal horn.” Journal of Neuroscience 24(11): 2774–2781.

Baurand, A., P. Raboisson, M. Freund, C. Léon, J.-P. Cazenave, J.-J. Bourguignon and C. Gachet (2001). “Inhibition of platelet function by administration of MRS2179, a P2Y1 receptor antagonist.” European journal of pharmacology 412(3): 213–221.

Bodin, P. and G. J. N. r. Burnstock (2001). “Purinergic signalling: ATP release.” 26(8-9): 959–969.

Bokil, H., N. Laaris, K. Blinder, M. Ennis and A. Keller (2001). “Ephaptic interactions in the mammalian olfactory system.” Journal of Neuroscience 21(20): RC173.

Bruzzone, R., M. T. Barbe, N. J. Jakob and H. Monyer (2005). “Pharmacological properties of homomeric and heteromeric pannexin hemichannels expressed in Xenopus oocytes.” Journal of neurochemistry 92(5): 1033–1043.

Brändle, U., P. Spielmanns, R. Osteroth, J. Sim, A. Surprenant, G. Buell, J. Ruppersberg, P. Plinkert, H.-P. Zenner and E. Glowatzki (1997). “Desensitization of the P2X2 receptor controlled by alternative splicing.” FEBS Letters 404(2-3): 294–298.

Burgard, E. C., W. Niforatos, T. van Biesen, K. J. Lynch, E. Touma, R. E. Metzger, E. A. Kowaluk and M. F. Jarvis (1999). “P2X receptor–mediated ionic currents in dorsal root ganglion neurons.” Journal of Neurophysiology 82(3): 1590–1598.

Carvalho, G. and A. Damasio (2019). “Non-Synaptic Transmission and the Foundations of Affect.”

Chen, Y., G. Li and L.-Y. M. Huang (2015). “p38 MAPK mediates glial P2X7R-neuronal P2Y1R inhibitory control of P2X3R expression in dorsal root ganglion neurons.” Molecular pain 11(1): 68.

Chen, Y., X. Zhang, C. Wang, G. Li, Y. Gu and L.-Y. M. Huang (2008). “Activation of P2X7 receptors in glial satellite cells reduces pain through downregulation of P2X3 receptors in nociceptive neurons.” Proceedings of the National Academy of Sciences USA 105(43): 16773–16778.

Coggeshall, R. E. and S. M. Carlton (1997). “Receptor localization in the mammalian dorsal horn and primary afferent neurons.” Brain Research Reviews 24(1): 28–66.

Damasio, A. and G. Carvalho (2013). “The nature of feelings: evolutionary and neurobiological origins.” Nature Reviews Neuroscience 14(2): 143–152.

Descarries, L., V. Gisiger and M. Steriade (1997). “Diffuse transmission by acetylcholine in the CNS.” Progress in Neurobiology 53(5): 603–625.

Descarries, L. and N. Mechawar (2000). Ultrastructural evidence for diffuse transmission by monoamine and acetylcholine neurons of the central nervous system. Progress in Brain Research, Elsevier. 125: 27–47.

Devor, M. (1999). “Unexplained peculiarities of the dorsal root ganglion.” Pain 82: S27–S35.

Devor, M. and P. D. Wall (1990). “Cross-excitation in dorsal root ganglia of nerve-injured and intact rats.” Journal of Neurophysiology 64(6): 1733–1746.

Du, X., H. Hao, S. Gigout, D. Huang, Y. Yang, L. Li, C. Wang, D. Sundt, D. B. Jaffe and H. Zhang (2014). “Control of somatic membrane potential in nociceptive neurons and its implications for peripheral nociceptive transmission.” Pain 155(11): 2306–2322.

Du, X., H. Hao, Y. Yang, S. Huang, C. Wang, S. Gigout, R. Ramli, X. Li, E. Jaworska and I. Edwards (2017). “Local GABAergic signaling within sensory ganglia controls peripheral nociceptive transmission.” The Journal of Clinical Investigation 127(5): 1741–1756.

Filippov, A. K., D. A. Brown and E. A. Barnard (2000). “The P2Y1 receptor closes the N-type Ca2+ channel in neurones, with both adenosine triphosphates and diphosphates as potent agonists.” British journal of pharmacology 129(6): 1063–1066.

Filippov, A. K., R. C. Choi, J. Simon, E. A. Barnard and D. A. Brown (2006). “Activation of P2Y1 nucleotide receptors induces inhibition of the M-type K+ current in rat hippocampal pyramidal neurons.” Journal of Neuroscience 26(36): 9340–9348.

Gafar, H., M. D. Rodriguez, G. K. Chandaka, I. Salzer, S. Boehm and K. Schicker (2016). “Membrane coordination of receptors and channels mediating the inhibition of neuronal ion currents by ADP.” Purinergic Signalling 12(3): 497–507.

Gerevich, Z., C. Müller and P. Illes (2005). “Metabotropic P2Y1 receptors inhibit P2X3 receptor-channels in rat dorsal root ganglion neurons.” European journal of pharmacology 521(1-3): 34–38.

Grasa, L., V. Gil, D. Gallego, M. T. Martín and M. Jiménez (2009). “P2Y1 receptors mediate inhibitory neuromuscular transmission in the rat colon.” British journal of pharmacology 158(6): 1641–1652.

Grubb, B. D. and R. J. Evans (1999). “Characterization of cultured dorsal root ganglion neuron P2X receptors.” European Journal of Neuroscience 11(1): 149–154.

Hechler, B., C. Nonne, E. J. Roh, M. Cattaneo, J.-P. Cazenave, F. Lanza, K. A. Jacobson and C. Gachet (2006). “MRS2500 [2-iodo-N6-methyl-(N)-methanocarba-2’-deoxyadenosine-3’, 5’-bisphosphate], a potent, selective, and stable antagonist of the platelet P2Y1 receptor with strong antithrombotic activity in mice.” Journal of Pharmacology and Experimental Therapeutics 316(2): 556–563.

Herkenham, M. (1987). “Mismatches between neurotransmitter and receptor localizations in brain: observations and implications.” Neuroscience 23(1): 1–38.

Hiruma, H. and C. Bourque (1995). “P2 purinoceptor-mediated depolarization of rat supraoptic neurosecretory cells in vitro.” The Journal of physiology 489(3): 805–811.

Huang, L. Y. M., Y. Gu and Y. Chen (2013). “Communication between neuronal somata and satellite glial cells in sensory ganglia.” Glia 61(10): 1571–1581.

Huettner, J. E., G. A. Kerchner and M. Zhuo (2002). “Glutamate and the presynaptic control of spinal sensory transmission.” The Neuroscientist 8(2): 89–92.

Jankowski, M. P., K. K. Rau, D. J. Soneji, K. M. Ekmann, C. E. Anderson, D. C. Molliver and H. R. Koerber (2012). “Purinergic receptor P2Y1 regulates polymodal C-fiber thermal thresholds and sensory neuron phenotypic switching during peripheral inflammation.” Pain 153(2): 410–419.

Jiménez, M., P. Clavé, A. Accarino and D. Gallego (2014). “Purinergic neuromuscular transmission in the gastrointestinal tract; functional basis for future clinical and pharmacological studies.” British journal of pharmacology 171(19): 4360–4375.

Kaiser, R. A. and I. L. Buxton (2002). “Nucleotide-mediated relaxation in guinea-pig aorta: selective inhibition by MRS2179.” British journal of pharmacology 135(2): 537–545.

Kim, H. S., M. Ohno, B. Xu, H. O. Kim, Y. Choi, X. D. Ji, S. Maddileti, V. E. Marquez, T. K. Harden and K. A. Jacobson (2003). “2-Substitution of adenine nucleotide analogues containing a bicyclo [3.1. 0] hexane ring system locked in a northern conformation: enhanced potency as P2Y1 receptor antagonists.” Journal of Medicinal Chemistry 46(23): 4974–4987.

Kim, Y. S., M. Anderson, K. Park, Q. Zheng, A. Agarwal, C. Gong, L. Young, S. He, P. C. LaVinka and F. Zhou (2016). “Coupled activation of primary sensory neurons contributes to chronic pain.” Neuron 91(5): 1085–1096.

King, C. H. and S. S. Scherer (2012). “Kv7. 5 is the primary Kv7 subunit expressed in C-fibers.” Journal of Comparative Neurology 520(9): 1940–1950.

Krames, E. S. (2014). “The role of the dorsal root ganglion in the development of neuropathic pain.” Pain Medicine 15(10): 1669–1685.

Lee, C. J., R. Bardoni, C.-K. Tong, H. S. Engelman, D. J. Joseph, P. C. Magherini and A. B. MacDermott (2002). “Functional expression of AMPA receptors on central terminals of rat dorsal root ganglion neurons and presynaptic inhibition of glutamate release.” Neuron 35(1): 135–146.

Li, J., J. A. McRoberts, J. Nie, H. S. Ennes and E. A. Mayer (2004). “Electrophysiological characterization of N-methyl-D-aspartate receptors in rat dorsal root ganglia neurons.” Pain 109(3): 443–452.

Lieberman, A. (1976). Sensory ganglia. In The Peripheral Nerve (edited by Landon, DN) pp. 188–278, London: Chapman and Hall. Google Scholar.

Locovei, S., J. Wang and G. Dahl (2006). “Activation of pannexin 1 channels by ATP through P2Y receptors and by cytoplasmic calcium.” FEBS letters 580(1): 239–244.

Magni, G. and S. Ceruti (2013). “P2Y purinergic receptors: new targets for analgesic and antimigraine drugs.” Biochemical pharmacology 85(4): 466–477.

Magni, G., D. Merli, C. Verderio, M. P. Abbracchio and S. Ceruti (2015). “P2Y2 receptor antagonists as anti-allodynic agents in acute and subchronic trigeminal sensitization: Role of satellite glial cells.” Glia 63(7): 1256–1269.

Malin, S. A. and D. C. Molliver (2010). “Gi-and Gq-coupled ADP (P2Y) receptors act in opposition to modulate nociceptive signaling and inflammatory pain behavior.” Molecular Pain 6(1): 21.

Matsuka, Y., J. K. Neubert, N. T. Maidment and I. J. B. r. Spigelman (2001). “Concurrent release of ATP and substance P within guinea pig trigeminal ganglia in vivo.” Brain Research 915(2): 248–255.

Matsuka, Y., T. Ono, H. Iwase, S. Mitrirattanakul, K. S. Omoto, T. Cho, Y. Y. N. Lam, B. Snyder and I. Spigelman (2008). “Altered ATP release and metabolism in dorsal root ganglia of neuropathic rats.” Molecular Pain 4(1): 66.

Matsuka, Y. and I. Spigelman (2004). “Hyperosmolar solutions selectively block action potentials in rat myelinated sensory fibers: implications for diabetic neuropathy.” Journal of Neurophysiology 91(1): 48–56.

Molliver, D. C., K. K. Rau, S. L. McIlwrath, M. P. Jankowski and R. Koerber (2011). “The ADP receptor P2Y 1 is necessary for normal thermal sensitivity in cutaneous polymodal nociceptors.” Molecular Pain 7(1): 13.

Oh, E. J. and D. Weinreich (2002). “Chemical communication between vagal afferent somata in nodose Ganglia of the rat and the Guinea pig in vitro.” Journal of Neurophysiology 87(6): 2801–2807.

Rajasekhar, P., D. P. Poole, W. Liedtke, N. W. Bunnett and N. A. Veldhuis (2015). “P2Y1 receptor activation of the TRPV4 ion channel enhances purinergic signaling in satellite glial cells.” Journal of Biological Chemistry 290(48): 29051–29062.

Ren, J., C. Qin, F. Hu, J. Tan, L. Qiu, S. Zhao, G. Feng and M. J. N. Luo (2011). “Habenula “cholinergic” neurons corelease glutamate and acetylcholine and activate postsynaptic neurons via distinct transmission modes.” Neuron 69(3): 445–452.

Rozanski, G. M., Q. Li and E. F. Stanley (2013). “Transglial transmission at the dorsal root ganglion sandwich synapse: glial cell to postsynaptic neuron communication.” European Journal of Neuroscience 37(8): 1221–1228.

Sawynok, J. and X. J. Liu (2003). “Adenosine in the spinal cord and periphery: release and regulation of pain.” Progress in neurobiology 69(5): 313–340.

Shigemoto, R., H. Ohishi, S. Nakanishi and N. Mizuno (1992). “Expression of the mRNA for the rat NMDA receptor (NMDAR1) in the sensory and autonomic ganglion neurons.” Neuroscience Letters 144(1-2): 229–232.

Shinder, V. and M. Devor (1994). “Structural basis of neuron-to-neuron cross-excitation in dorsal root ganglia.” Journal of Neurocytology 23(9): 515–531.

Sokolova, E., A. Skorinkin, I. Moiseev, A. Agrachev, A. Nistri and R. Giniatullin (2006). “Experimental and modeling studies of desensitization of P2X3 receptors.” Molecular Pharmacology.

Spigelman, I., Z. Li, P. K. Banerjee, R. M. Mihalek, G. E. Homanics and R. W. Olsen (2002). “Behavior and physiology of mice lacking the GABAA-receptor δ subunit.” Epilepsia 43: 3–8.

Sykova, E. (2004). “Extrasynaptic volume transmission and diffusion parameters of the extracellular space.” Neuroscience 129(4): 861–876.

Sykova, E. (2005). “Glia and volume transmission during physiological and pathological states.” Journal of Neural Transmission 112(1): 137–147.

Syková, E. and A. Chvátal (2000). “Glial cells and volume transmission in the CNS.” Neurochemistry International 36(4-5): 397–409.

Vizi, E. S., A. Fekete, R. Karoly and A. Mike (2010). “Non-synaptic receptors and transporters involved in brain functions and targets of drug treatment.” British Journal of Pharmacology 160(4): 785–809.

Vongtau, H., E. Lavoie, J. Sévigny and D. J. N. Molliver (2011). “Distribution of ectonucleotidases in mouse sensory circuits suggests roles for nucleoside triphosphate diphosphohydrolase-3 in nociception and mechanoreception.” Pain Mechanisms and Sensory Neuroscience 193: 387–398.

Warwick, R. A. and M. Hanani (2016). “Involvement of aberrant calcium signalling in herpetic neuralgia.” Experimental neurology 277: 10–18.

Xu, G.-Y. and L.-Y. M. Huang (2002). “Peripheral inflammation sensitizes P2X receptor-mediated responses in rat dorsal root ganglion neurons.” Journal of Neuroscience 22(1): 93–102.

Xu, G.-Y. and Z.-Q. Zhao (2003). “Cross-inhibition of mechanoreceptive inputs in dorsal root ganglia of peripheral inflammatory cats.” Brain research 970(1-2): 188–194.

Yousuf, A., F. Klinger, K. Schicker and S. Boehm (2011). “Nucleotides control the excitability of sensory neurons via two P2Y receptors and a bifurcated signaling cascade.” Pain 152(8): 1899–1908.

Zhang, D., Z.-G. Gao, K. Zhang, E. Kiselev, S. Crane, J. Wang, S. Paoletta, C. Yi, L. Ma and W. Zhang (2015). “Two disparate ligand-binding sites in the human P2Y 1 receptor.” Nature 520(7547): 317.

Zhang, Y., G. Laumet, S.-R. Chen, W. N. Hittelman and H.-L. Pan (2015). “Pannexin-1 up-regulation in the dorsal root ganglion contributes to neuropathic pain development.” Journal of Biological Chemistry 290(23): 14647–14655.

